# Unraveling the Pathogenesis of Idiopathic Pulmonary Fibrosis Complicated with Type 2 Diabetes through Microarray Data Analysis

**DOI:** 10.1101/2023.07.31.551395

**Authors:** Zihan Xu

## Abstract

**Background:** Idiopathic pulmonary fibrosis (IPF) is a chronic and progressive lung disease characterized by excessive scarring of lung tissue. Recent studies have indicated a potential link between IPF and type 2 diabetes (T2D), suggesting that T2D may contribute to the pathogenesis of IPF or vice versa. In this study, we aim to investigate the underlying molecular mechanisms and pathways involved in the development of IPF complicated with T2D using microarray data analysis.

**Methods:** The datasets for Type 2 Diabetes (T2D) (GSE25724) and Idiopathic Pulmonary Fibrosis (IPF) (GSE110147) were obtained from the Gene Expression Omnibus (GEO) database. Differential expression analysis was conducted using the ‘limma’ package in R to identify genes that were significantly differentially expressed between T2D and IPF samples. Functional enrichment analysis of these differentially expressed genes was performed using Gene Ontology and Kyoto Encyclopedia of Genes and Genomes (KEGG).

To explore protein-protein interactions, protein-protein interaction networks were constructed using the Search Tool for the Retrieval of Interacting Genes (STRING) database and visualized using Cytoscape. CytoHubba, a plugin in Cytoscape, was utilized to identify hub genes in the network. The identified hub genes were further validated in independent datasets: GSE166467 for T2D and GSE24206 for IPF.

To assess the predictive value of the hub genes, receiver operating characteristic (ROC) curves were generated. Additionally, gene set enrichment analysis was performed to uncover potential biological pathways associated with the hub genes. Finally, an analysis of immune infiltration within the hub gene network was conducted.

**Results:** A total of 255 differentially expressed genes (DEGs) were found to be commonly dysregulated in both Type 2 Diabetes (T2D) and Idiopathic Pulmonary Fibrosis (IPF). Pathways related to metabolic processes were significantly enriched in the analysis of both T2D and IPF. Through the validation process, RPL30 was identified as a hub gene, exhibiting an area under the curve (AUC) value greater than 0.65 for both T2D and IPF. Additionally, we identified 92 transcription factors (TFs) and 54 microRNAs (miRNAs) that may potentially regulate the expression of RPL30.

**Conclusions:** This study provides novel insights by highlighting the role of RPL30 in the shared pathogenesis of pulmonary fibrosis and Type 2 diabetes (T2D). The findings suggest that RPL30 has the potential to serve as a biomarker and therapeutic target for these conditions.

## Introduction

Diabetes is clinically categorized into two primary types: type 1 diabetes mellitus (T1D) and type 2 diabetes mellitus (T2D). Diabetes mellitus is a systemic metabolic disorder characterized by insulin deficiency or resistance, leading to consistently elevated blood glucose levels [1–4]. T2D, being the most prevalent form of diabetes, accounts for approximately 90% of all cases [5]. Prolonged hyperglycemia associated with diabetes can lead to micro- and macro-vascular complications, particularly affecting the renal, retinal, nervous, and cardiovascular systems [6]. According to the latest report by the International Diabetes Federation (IDF), the global population of adult diabetes mellitus patients is projected to reach 530 million, with an estimated increase to 780 million by 2045 [7]. Consequently, diabetes not only causes significant individual suffering but also imposes substantial clinical and financial burdens.

Idiopathic pulmonary fibrosis (IPF) is a chronic and diffuse interstitial lung disease of unknown origin. This disease predominantly affects the elderly population, with incidence rates rising in correlation with age [8,9]. A comprehensive analysis of global IPF incidence and mortality rates revealed that the incidence of IPF is comparable to that of liver, stomach, and cervical cancers, and the mortality rate of IPF is progressively increasing in numerous countries [10].

However, although the researches and evidences about lung is the target organ of IPF are lacking, hyperglycemia has been shown to lead to interstitial fibrosis and alveolar capillary microvascular pathologies [11,12]. These findings indicate that the lung may be susceptible to diabetes-related complications, highlighting its potential as a target organ. Furthermore, several clinical studies have suggested a potential association between diabetes mellitus (DM) and idiopathic pulmonary fibrosis (IPF). A recent study conducted in Japan revealed a high prevalence of DM among IPF patients, thus identifying DM as a significant risk factor for the development of IPF [13]. A nationwide survey conducted in South Korea revealed that patients with idiopathic pulmonary fibrosis (IPF) exhibit higher rates of type 2 diabetes (T2D) compared to a matched control population. Additionally, IPF patients with diabetes show a significantly higher prevalence of hypertension, cardiovascular disease, and other malignant tumors when compared to IPF patients without diabetes [14]. Furthermore, it has been observed that IPF patients with diabetes tend to have longer hospital stays, implying a poorer prognosis [15,16]. Despite strong clinical and epidemiological evidence supporting the association between IPF and T2D [17–20], the causal relationship remains unclear, and little is known about the shared molecular signatures and gene regulatory mechanisms. Therefore, elucidating the key molecular mechanisms underlying the connection between IPF and T2D can not only enhance our understanding of the molecular pathogenesis of both diseases but also facilitate the development of novel therapeutic strategies.

In this study, we first analyzed two data sets downloaded from the GEO (https://www.ncbi.nlm.nih.gov/geo/) to obtain the shared genes of IPF and T2D. Then the co-DEGs were analyzed through GO, KEGG and PPI networks to clarify their biological functions and pathways. We also predicted the hub genes through various algorithms, and evaluated the diagnostic performance of the hub genes in the validation datasets. In addition, we predicted the potential transcription factors (TFs) and miRNAs of the hug genes and constructed a visual interaction network. Finally, we analyzed the relationship between hub genes and immune infiltration. The research results preliminarily infer the common biomarkers and pathways involved in the pathogenesis of IPF and T2D, which will help to clarify the common pathological mechanism of IPF and T2D.

In this study, we conducted an analysis using two publicly available data sets obtained from the Gene Expression Omnibus (GEO) database (https://www.ncbi.nlm.nih.gov/geo/) to identify the shared genes between idiopathic pulmonary fibrosis (IPF) and type 2 diabetes (T2D). Subsequently, we performed gene ontology (GO), Kyoto Encyclopedia of Genes and Genomes (KEGG), and protein-protein interaction (PPI) network analyses to elucidate the biological functions and pathways associated with the co-differentially expressed genes (co-DEGs). Additionally, we employed various algorithms to predict hub genes and evaluated their diagnostic performance using independent validation datasets. Moreover, we predicted potential transcription factors (TFs) and microRNAs (miRNAs) that may regulate the identified hub genes and constructed a visual interaction network. Lastly, we analyzed the relationship between the hub genes and immune cell infiltration. Our findings provide preliminary insights into the shared biomarkers and pathways involved in the pathogenesis of both IPF and T2D, contributing to a better understanding of their common underlying mechanisms.

## Methods

### Data source

The datasets GSE25724 [21], GSE166467 [22], GSE110147 [23], and GSE24206 [24] were retrieved from the GEO database (http://www.ncbi.nlm.nih.gov/gds/). Table 1 provides detailed information on these datasets.

**Table 1.** Summary of those four GEO datasets involving T2D and IPF patients.

| ID | GSE number | Platform | Samples | Disease | Group |
| --- | --- | --- | --- | --- | --- |
| 1 | GSE25724 | GPL96 | 6 patients and 7 controls | T2D | Discovery |
| 2 | GSE166467 | GPL10558 | 13 patients and 13 controls | T2D | Validation |
| 3 | GSE110147 | GPL6244 | 22 patients and 11 controls | IPF | Discovery |
| 4 | GSE24206 | GPL570 | 17 patients and 6 controls | IPF | Validation |

### Identification of differentially expressed genes (DEGs)

We obtained the series matrix files of the datasets from GEO. For GSE25724 and GSE110147, the data were processed using the Limma package [25] in R to identify the differential expression of mRNAs between case and control samples. Genes with a significant differential expression in each dataset were defined as having an absolute log2 Fold Change (log FC) value greater than 1.0 and a Benjamini-Hochberg adjusted p-value lower than 0.05. The online Venn diagram tool available at http://bioinformatics.psb.ugent.be/webtools/ was utilized to identify the common DEGs.

### Functional analysis

To elucidate the functions of the common DEGs, we utilized the GO database (http://geneontology.org/) and KEGG database (https://www.genome.jp/kegg/) to obtain in-depth information on biological processes and signaling pathways. A P-value < 0.05 was employed as a standardized measure to quantify the top functional categories and pathways. The GO database, established by the Gene Ontology Federation, provides concise annotations of gene products, including their functions, involvement in biological pathways, and cellular location [26]. On the other hand, the KEGG database is dedicated to storing information on gene pathways across various species [27]. Additionally, we also incorporated the WikiPathways, Reactome, and BioCarta databases to gain a more comprehensive understanding of the pertinent signaling pathways.

### Protein–protein interaction Network Construction and Module Analysis

As for the construction of the protein-protein interaction (PPI) network, the STRING database (https://string-db.org/) was utilized, with a minimum interaction score of 0.4 considered as the confidence threshold. Moreover, all other parameters were set to their default values. Subsequently, the MCODE plug-in implemented in Cytoscape was employed to identify the significant functional modules, with the default parameters utilized for this analysis.

### Hub Genes Selection and Validation of the Expression

We then used cytoHubba to identify the hug genes. Here, six common algorithms (MCC, MNC, Degree, EPC, EcCentricity and Closeness) were used to evaluate and select hub genes. The expression levels of the identified hub genes were validated in GSE166467 for T2D and GSE24206 for IPF. The comparison between the two sets of data was performed with the T-test. P-value < 0.05 was considered significant. Subsequently, GeneMANIA (http://www.genemania.org/) was used to construct a co-expression network of hub gene.

### Receiver Operating Characteristic Curves and Gene-regulatory network analysis

In accordance with the conventions of SCI paper formatting, we evaluated the predictive accuracy of the hub genes through the generation of Receiver Operating Characteristic (ROC) curves. This was achieved by utilizing the pROC package [28] in the R language.

To gain insights into the interaction between common genes and transcription factors, as well as to assess the impact of these factors on the expression and functional pathways of such genes, we employed the JASPAR database [29], ChEA database [30], and ENCODE database [31].

Furthermore, DEG-miRNA interaction networks were discovered by leveraging the TarBase and miRTarBase databases, facilitated through the usage of NetworkAnalyst (https://www.networkanalyst.ca/). The obtained outcomes were subsequently visualized using Cytoscape.

### candidate drugs prediction

To predict protein-drug interactions and identify potential candidate drugs, we utilized the DSigDB database (http://dsigdb.tanlab.org/DSigDBv1.0/) by uploading the common genes. Significance was determined based on a P-value threshold of less than 0.05.

### Gene–disease-association analysis

In order to enhance our comprehension of human genetic disorders, we leveraged DisGeNET (http://www.disgenet.org/). This resource is dedicated to expanding our knowledge of such disorders. Additionally, we employed NetworkAnalyst to explore gene-disease associations, aiming to unveil diseases that are linked to the common DEGs.

## Results

### Identification of shared transcriptomic signature between T2D and IPF

Figure 1 depicts the crucial procedures used in this study. Four GEO datasets - GSE25724, GSE166467, GSE110147, and GSE24206 - were selected for this analysis. Table 1 summarizes the information of these datasets.

**Figure 1.**
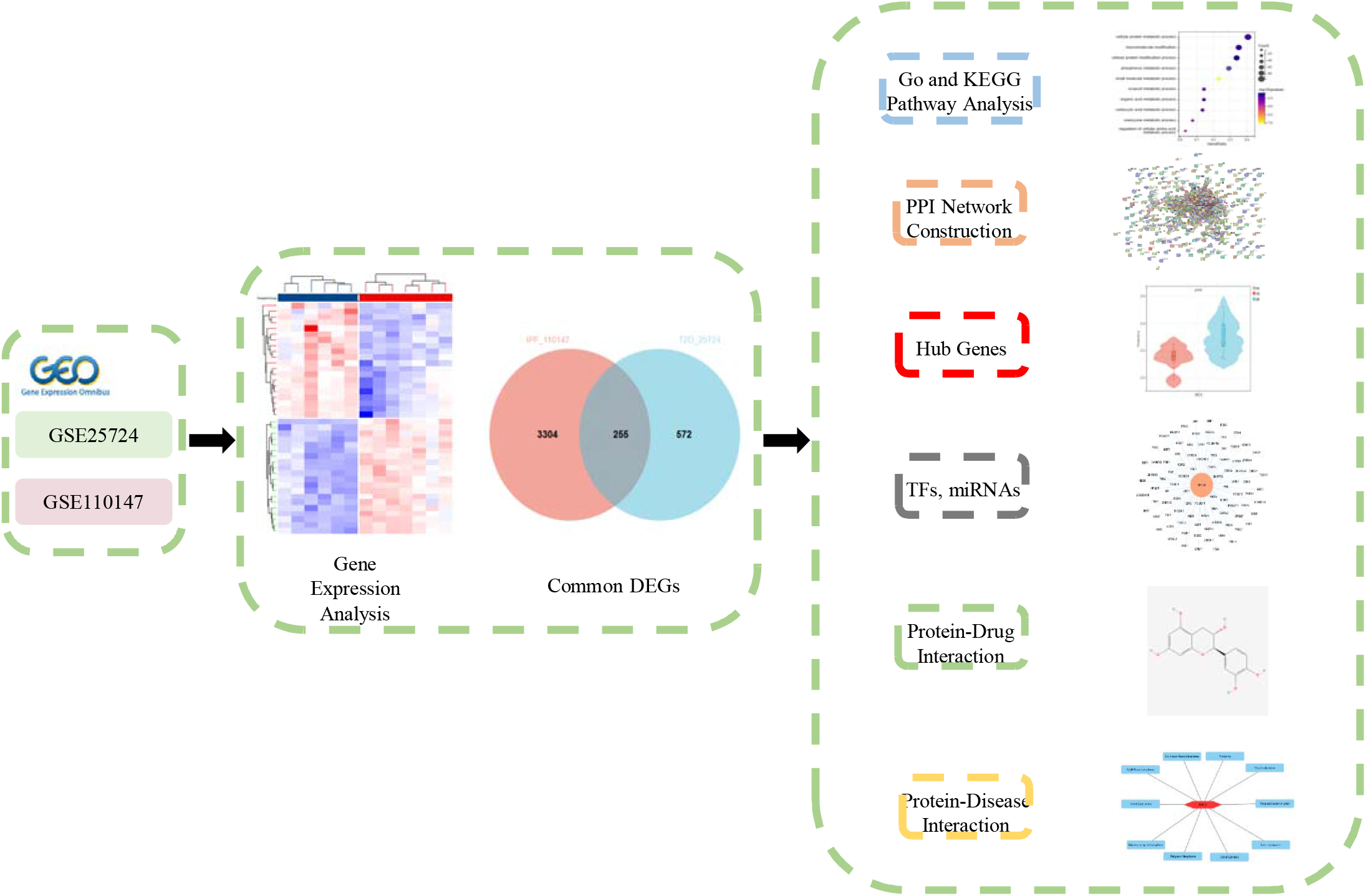
The predominant methodical workflow of this study.

In the DEG analysis, GSE25724 and GSE110147 were combined as discovery cohorts, while GSE166467 and GSE24206 were paired as validation cohorts. Using a cutoff criteria of absolute log2FC > 1 and adjusted P value < 0.05, differentially expressed genes (DEGs) were extracted from GSE25724 and GSE110147 after standardizing the microarray results. In T2D patients compared to normal individuals, 827 DEGs were identified, while in IPF patients compared to healthy controls, a total of 3559 DEGs were found.

For better visualization, volcano plots were used to present the dysregulated genes. Upregulated genes were represented by red dots, while downregulated genes were represented by green dots. Figure 2A shows the volcano plot for T2D, and Figure 2B shows the one for IPF. By taking the intersection of the Venn diagram, 255 shared DEGs were obtained and integrated into the downstream analysis, as shown in Figure 2C.

**Figure 2.**
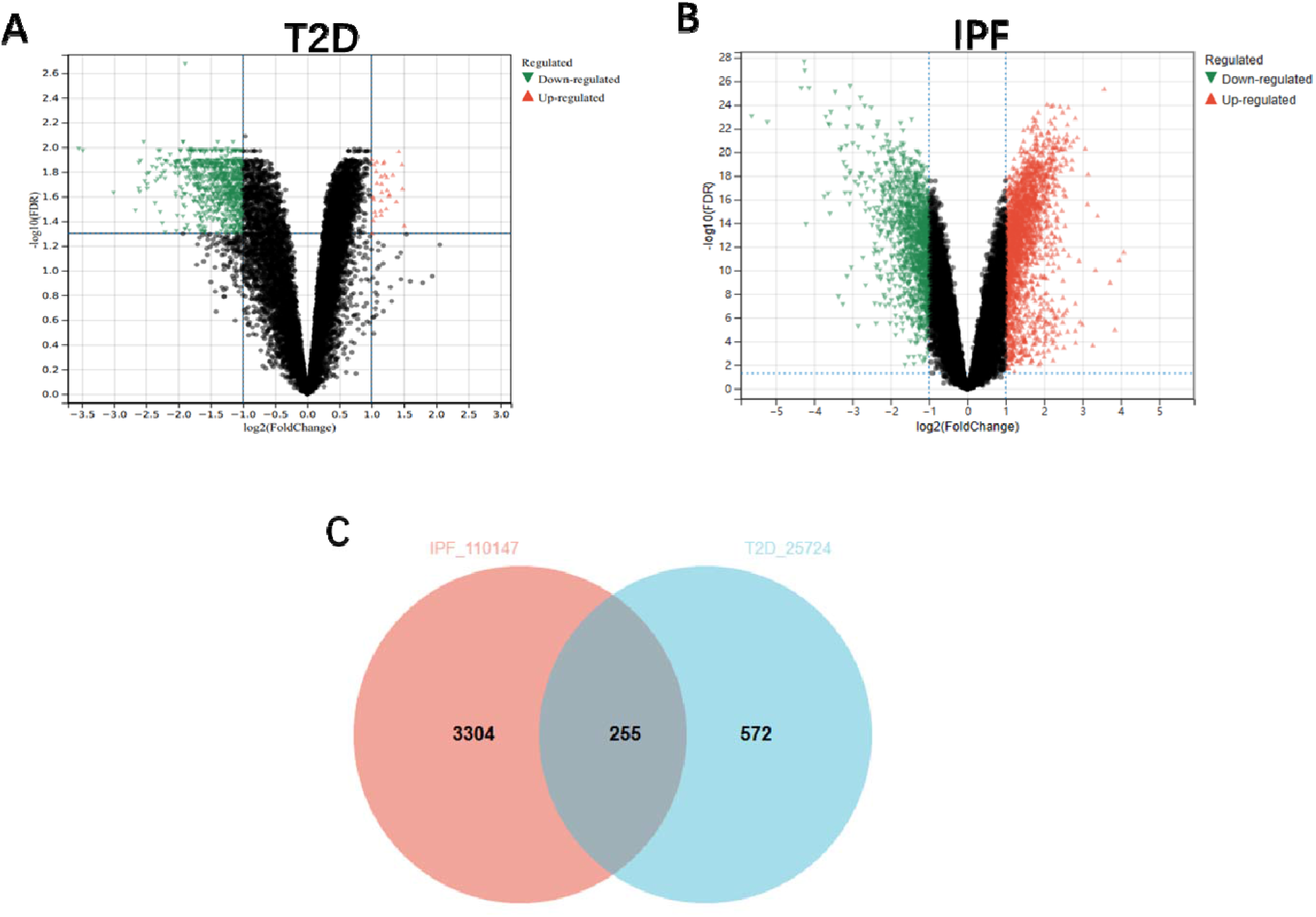
Identification of common DEGs. DEGs in GSE25724 (A) and GSE110147 (B) were shown in the volcano plot, with red colour representing significantly high expression genes and green dots representing significantly down-regulated genes. (C) Venn diagram depicted concordant genes from the intersection of two datasets GSE25724 and GSE110147.

### Analysis of the Functional Characteristics of Common DEGs

To further explore the underlying biological information of these common DEGs, the set of databases (GO, KEGG, Reactome, WikiPathways, and BioCarta) was utilized to identify enriched functional terms.

Gene ontology, pathway enrichment analysis, and the visualization of results were executed by Enrichr. Using the GO database as an annotation source, the GO analysis of common genes was divided into three categories (biological process, cellular component, and molecular function). The GO terms for each of the three predominant categories were collected in Supplementary File 2 and presented in the form of dot charts in Figure 3. The differentially expressed genes were significantly enriched in small molecule metabolic process, cytosol, and nucleotide binding in the biological processes, cell compartments, and molecular functions subsets, respectively.

**Figure 3.**
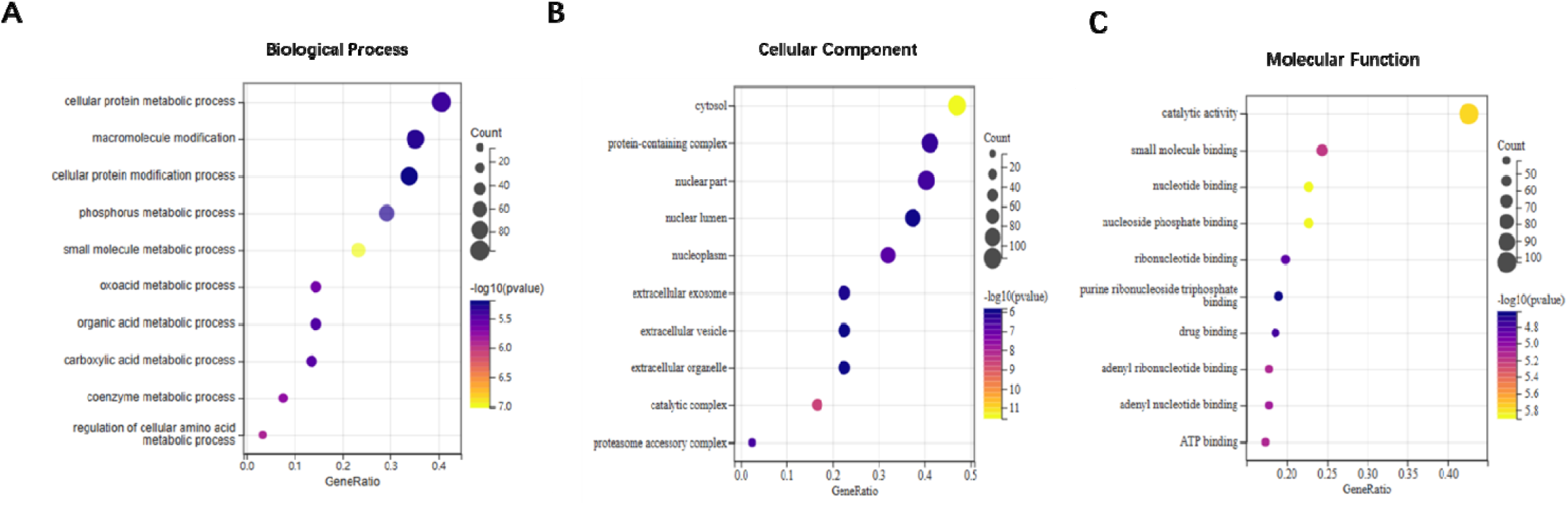
GO terms of common genes between T2D and IPF. (A)Biological Processes, (B) Cellular Component, (C) Molecular Function.

The most impacted pathways of the common genes among T2D and IPF were gathered from four global databases, including KEGG, BioCarta, Reactome and WikiPathway. KEGG pathway analysis revealed the following top 10 pathways: Proteasome, Epstein-Barr virus infection, Citrate cycle (TCA cycle), Metabolic pathways, Amino sugar and nucleotide sugar metabolism, Cysteine and methionine metabolism, Sulfur metabolism, Propanoate metabolism, Influenza A and Phosphatidylinositol signaling system. The pathways found in the examined datasets are listed in Supplementary File 3 and presented in the form of pie charts in Figure 4.

**Figure 4.**
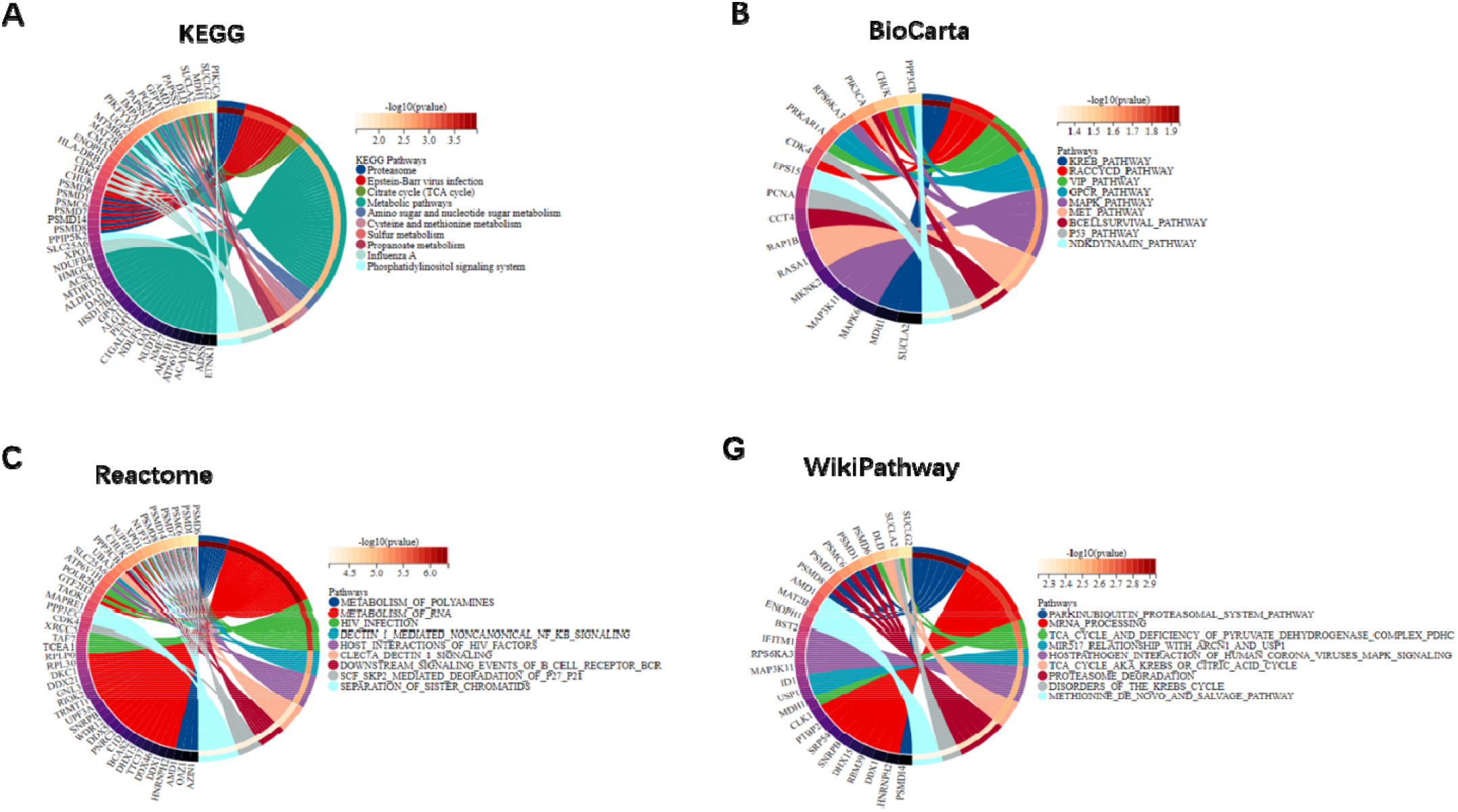
Pathway enrichment analysis of common genes between T2D and IPF. (A) KEGG Human Pathway, (B) BioCarta Pathway, (C) Reactome Pathway, (D) Wikipathway.

### PPI Network Construction of Common DEGs

To benefit clinically and pharmacologically oriented research for further biological studies, the 255 common genes were uploaded to the STRING database. As shown in Figure 5A, the PPI network of common genes consists of 253 nodes and 647 edges. MCODE, a plug-in of Cytoscape, was used to conduct module analysis to detect crucial clustering modules. Seven closely connected gene modules were obtained. Module 1 included 18 nodes and 68 edges with a cluster score of 8 (Figure 5B).

**Figure 5.**
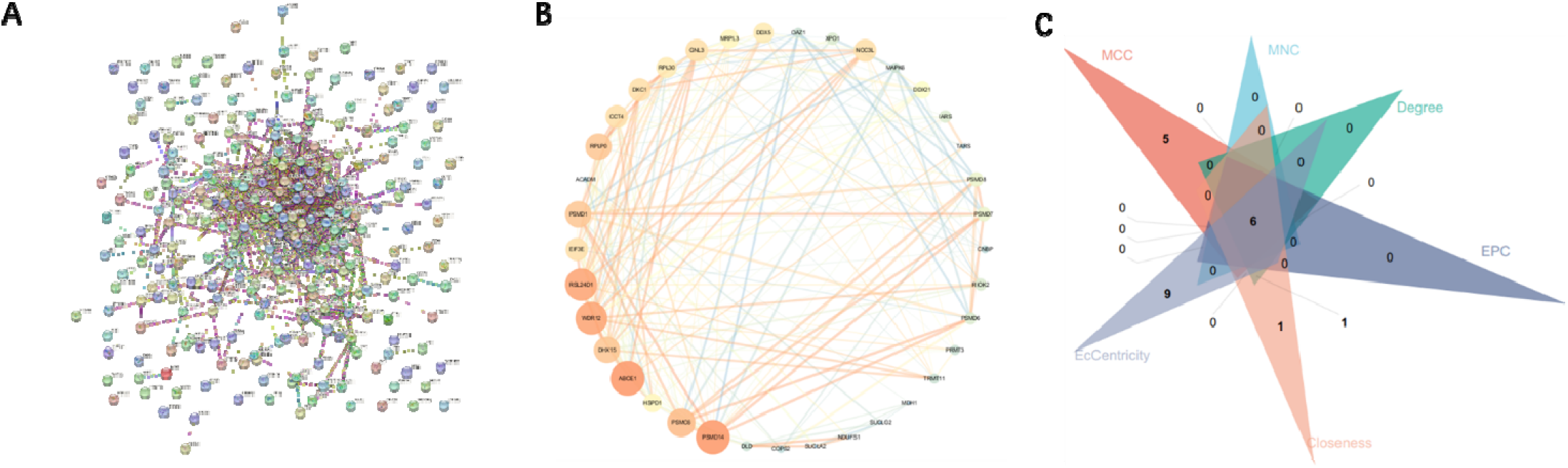
PPI Network Construction of Common DEGs. (A) PPI network of the common DEGs constructed by STRING. (B) Significant gene module and enrichment analysis of the modular genes. (C) Identification of 6 candidates for hub genes by five algorithms.

### Identification and Validation of Hub Genes

To explore genes that may play an important role in the co-occurrence of T2D and IPF, we have calculated the top 20 hub genes through the six algorithms of plug-in cytoHubba (Table 2). The intersection of these 20 genes from the six algorithms revealed 6 candidate hub genes: ABCE1, PSMD14, PSMD1, PSMC6, RPL30 and RPLP0 (Figure 5C).

**Table 2.** The top 20 hub genes rank in cytoHubba.

| MCC | MNC | Degree | EPC | Closeness | EcCentricity |
| --- | --- | --- | --- | --- | --- |
| WDR12 | PSMD14 | PSMC6 | PSMD14 | PSMC6 | XPO1 |
| RSL24D1 | DHX15 | DHX15 | DHX15 | PSMD14 | OAZ1 |
| NOC3L | ABCE1 | PSMD14 | ABCE1 | DHX15 | RPLP0 |
| DHX15 | PSMC6 | ABCE1 | PSMC6 | ABCE1 | HSPD1 |
| GNL3 | CCT4 | XPO1 | WDR12 | XPO1 | RPL30 |
| DKC1 | XPO1 | CCT4 | RSL24D1 | CCT4 | PSMD14 |
| DDX5 | WDR12 | WDR12 | RPLP0 | DDX1 | CCT4 |
| DDX21 | RSL24D1 | RSL24D1 | CCT4 | WDR12 | PSMD1 |
| ABCE1 | RPLP0 | DDX5 | DDX5 | RSL24D1 | IARS |
| RIOK2 | DDX5 | RPLP0 | PSMD1 | DDX5 | EIF3E |
| PSMD14 | GNL3 | DDX1 | DDX21 | RPLP0 | ABCE1 |
| TRMT11 | RPL30 | RPL30 | NOC3L | PSMD1 | PSMC6 |
| PSMD1 | DDX21 | PSMD1 | DDX1 | HSPD1 | SLC25A6 |
| PSMC6 | PSMD1 | GNL3 | GNL3 | IARS | USP16 |
| RPLP0 | DKC1 | DDX21 | DKC1 | PCNA | RNF138 |
| EIF3E | MRPL3 | DKC1 | XPO1 | DDX21 | PAPSS2 |
| PSMD7 | HSPD1 | IARS | EIF3E | EIF3E | PAPSS1 |
| PSMD8 | DDX1 | PCNA | MRPL3 | MRPL3 | PDCD10 |
| PSMD6 | IARS | HSPD1 | RPL30 | MATR3 | MAPK6 |
| RPL30 | EIF3E | NOC3L | IARS | RPL30 | ZFAND1 |

In order to verify the reliability of these hub gene expression levels, validation was performed in GSE166467 for T2D and GSE24206 for IPF from GEO. The results showed that only RPL30 was significantly upregulated in both T2D and IPF compared with normal tissues (Figure 6A&D). Moreover, the expression level of the other 5 candidate genes in validation datasets were shown in Supplementary Figure 1. Meanwhile, spearman correlation matrix analysis was also performed to identify the hub gene pairs with obvious correlation at the expression levels in T2D (Figure 6B) and IPF (Figure 6E).

**Figure 6.**
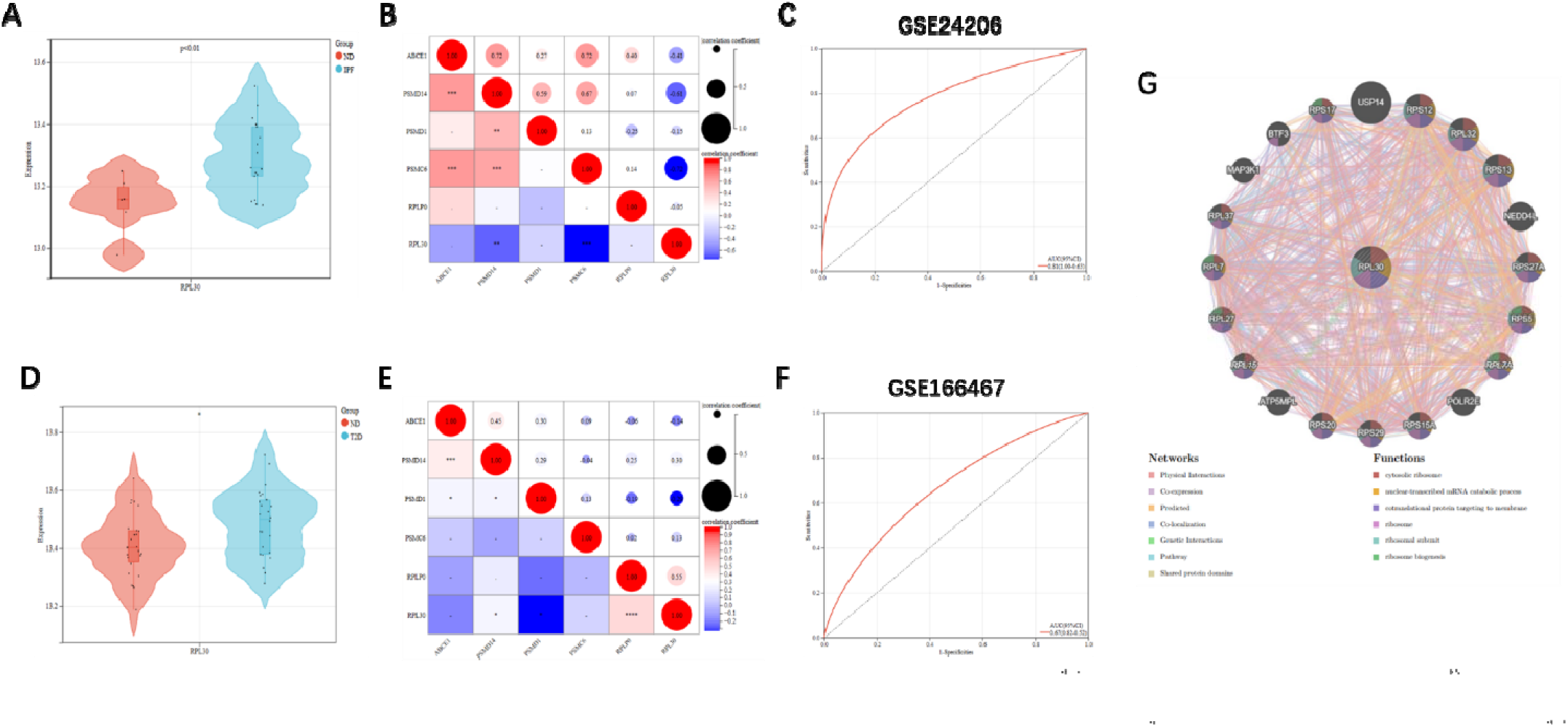
Validation of hub gene. (A) RPL30 was validated in GSE24206. (B) Spearman correlation matrix evaluated on the gene expression profile of hub genes in T2D. (C) ROC curves were drawn to evaluate the accuracy of the RPL30 in diagnosing T2D. (D) RPL30 was validated in GSE166467. (E) Spearman correlation matrix evaluated on the gene expression profile of hub genes in IPF. (F) ROC curves were drawn to evaluate the accuracy of the RPL30 in diagnosing T2D. (G) RPL30 and its co-expression genes was analyzed via GeneMANIA. *p < 0.05, **p < 0.01, ***p < 0.001.

### Efficacy Evaluation and PPI Construction of Hub Genes

To verify the diagnostic performance of the shared hub genes, ROC curves were drawn in the external T2D and IPF datasets. The AUC value in ROC curve demonstrated that the high accuracy of predictive value in the hub gene (Figures 6C&F). These results indicated that RPL30 can be promising markers for diagnosing T2D and IPF.

Based on GeneMANIA’s functional annotation patterns, a co-expression PPI network of RPL30 and its 20 interaction genes was established to predict the related functions and pathways. The predicted genes are located in the outer circle, while hub genes are in the inner circle. As shown in Figure 6G, the network illustrates that these genes are enriched in cytosolic ribosome, nuclear-transcribed mRNA catabolic process, cotranslational protein targeting to membrane, ribosome, ribosomal subunit, ribosome biogenesis.

### GRN analysis identified DEG–miRNA and Transcription factor–gene interaction networks

In order to determine the major changes at the transcriptional level and better understand the regulatory hub protein, Figure 7A depicts the interactions network between TFs and RPL30. The TF–gene interaction network consists of 93 nodes. DEG–miRNA coregulatory network is also generated (Figure 7B). The DEG–miRNA coregulatory network comprises 55 nodes.

**Figure 7.**
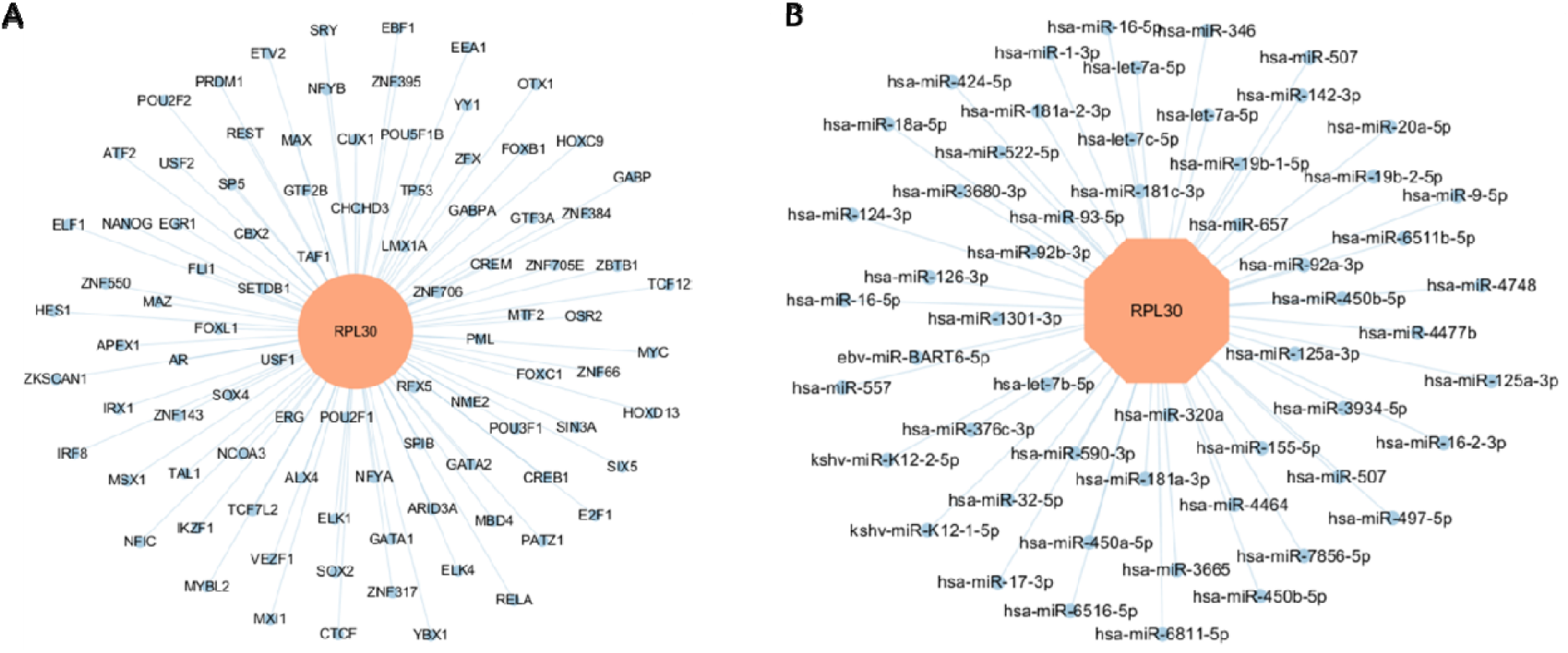
DEG–miRNA- and TF–gene-interaction networks. (A) Interconnected regulatory interaction network of differentially expressed genes (DEGs)–transcription factors (TFs) created using the Network Analyst. Herein, blue nodes represent TFs. (B) Interconnected regulatory interaction network of differentially expressed genes (DEGs)–miRNAs. Herein, the blue circle nodes represent miRNAs.

### Identification of candidate pharmacologic agents

To facilitate further research on therapy development, we made predictions about possible effective intervention drugs by using the DSigDB database. Table 3 presented the potential therapeutic drugs.

**Table 3.** Predictive top 10 pharmacologic agent candidates.

| Name | P-value | chemical formulate |
| --- | --- | --- |
| Beryllium sulfate CTD 00001005 | 0.001899964 | BeO4S |
| midecamycin HL60 DOWN | 0.011899889 | C41H67NO15 |
| testosterone enanthate CTD 00000155 | 0.026649833 | C26H40O3 |
| Cianidanol CTD 00005600 | 0.043499796 | C15H14O6 |

### Identification of disease associations

Different diseases can be related in some cases, such as when they share one or more similar genes. Decoding the relationship between genes and diseases is a crucial first step in in the development of treatment methods. Our investigation of gene–disease relationships with NetworkAnalyst (Figure 8) showed that Malignant Neoplasms, Primary malignant neoplasm, Colon Carcinoma, Carcinogenesis, Adult Medulloblastoma, Childhood Medulloblastoma, Myopathy, Medulloblastoma, Malignant tumor of colon and Liver carcinoma were most coordinated with the hub genes identified in this study.

**Figure 8.**
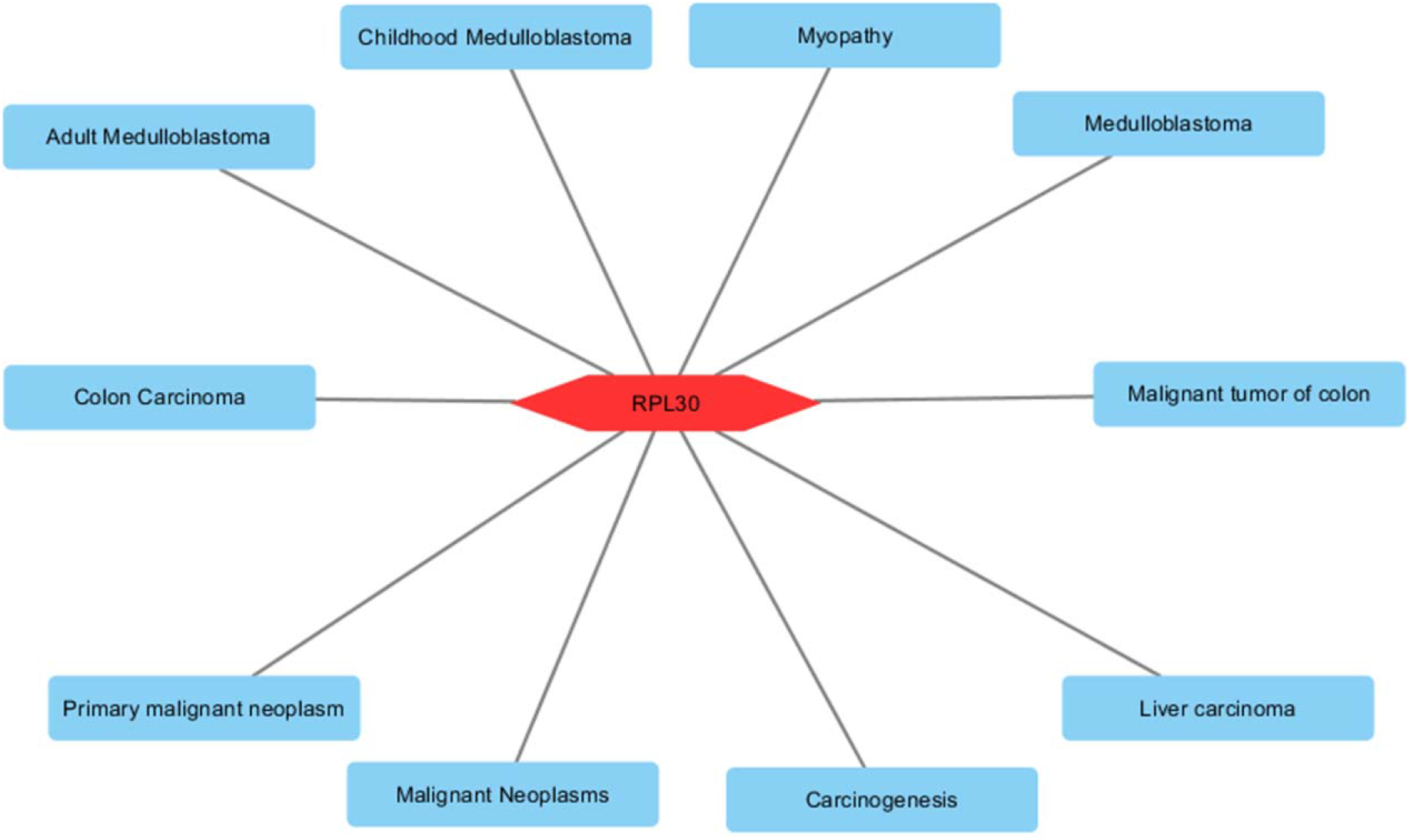
Gene-disease association network. Herein, the diseases represented by the blue nodes and the red nodes indicate the gene symbols that interact with the disease.

## Discussion

Both Type 2 Diabetes (T2D) and Idiopathic Pulmonary Fibrosis (IPF) are global epidemics that impose a significant burden on patients and healthcare systems. Increasing reports have confirmed the correlation between T2D and IPF [13,14,17,32,33]. A meta-analysis conducted previously demonstrated that IPF patients had a 1.54-fold increased odds of developing diabetes compared to non-IPF controls [20]. Additionally, another study revealed that the prevalence of diabetes in IPF patients was 25.4% compared to 13.4% in control subjects [34]. Furthermore, various studies have reported prevalence rates of diabetes in IPF patients ranging from 11.3% to 33% [13,35]. Moreover, several reports have suggested that T2D may act as an independent risk factor for IPF [13–20]. However, the precise underlying mechanism linking T2D and IPF remains incompletely understood. Hence, we conducted a series of bioinformatics analyses to explore the molecular mechanisms associated with T2D and IPF. Our goal is to investigate the potential pathophysiological relationship between IPF and T2D. In this study, we performed a series of bioinformatic analyzes on the T2D dataset GSE25724 and the IPF dataset GSE110147, and then identified 255 DEGs shared by T2D patients and IPF patients. The results of GO enrichment analysis showed that DEGs were mainly enriched in small molecule metabolic process, regulation of cellular amino acid metabolic process, coenzyme metabolic process, oxoacid metabolic process, carboxylic acid metabolic process, organic acid metabolic process. This result suggests that DEGs between T2D and IPF are closely related to metabolism in vivo. In addition, KEGG analysis showed that DEGs were mainly enriched in Citrate cycle (TCA cycle), Metabolic pathways, Amino sugar and nucleotide sugar metabolism, Cysteine and methionine metabolism, Sulfur metabolism, Propanoate metabolism. We further classified these pathways according to the KEGG database. We found that they are mainly related to metabolism. Many studies have shown that T2D and IPF are metabolic disorders. The results of functional enrichment of DEGs in this study are consistent with previous studies [36,37]. In addition, we built a PPI network based on the interaction between STRING database genes and shared DEGs. In order to further identify the hub genes, we then use the Cytoscape plug-in cytoHubba to score and identify the top 20 hub genes according to seven algorithms, such as MCC, MNC, Degree, EPC, Closeness and EcCentricity, and take the intersection. Then the expression level of these six shared hugs genes (ABCE1, PSMD14, PSMD1, PSMC6, RPLP0, RPL30) was verified using external data sets (GSE166467, GSE24206). Interestingly, we found that RPL30 was significantly up-regulated in T2D and IPF, suggesting that RPL30 might play an important role in the development of T2D and IPF.RPL30 gene encodes a ribosomal protein, which belongs to a member of ribosomal protein L30E family and is a component of 60S subunit [38]. Previous studies have shown that the related pathways of RPL30 are peptide chain elongation and rRNA processing in the nucleus and cytosol [39,40]. Recent clinical and animal studies have indicated that both insufficient and excessive Se intakes may increase risk of T2D, perhaps by way of selenoproteins actions [41–43]. Considering that RPL30 is involved in the regulation of selenium metabolism [44], we believe that RPL30 may cause T2D by interfering with selenium metabolism pathway. Another study reported that RPL30 may be involved in the dissociation of L13a and ribosomal 60s subunit, and free phosphorylated L13a may mediate the translation silencing of ceruloplasmin expression [45]. Ceruloplasmin can resist the ROS released by inflammatory cells [46]. During fibrosis, excessive oxidative stress will damage lung epithelial cells, and then induce lung fibrosis [47]. In addition, RPL30 has also been identified as a potential biomarker or therapeutic target for depression [48] and schizophrenia [49]. A recent study has shown that the hepatocarcinogenesis through NAFLD&NASH was induced through DNA methylation of RPL30 [50]. De Bortoli M and his colleagues indicated that the expression levels of RPL30 were adversely associated with overall and progression-free survival in medulloblastoma [51]. However, to the best of our knowledge, there is still no study so far that directly correlates the expression of RPL30 gene with T2D or IPF disease, so it is necessary to study the role of RPL30 in T2D and IPF pathology in the future.

TFs and miRNAs both play an important role in regulating gene expression. TFs are the key determinant of transcription of various target genes and many human diseases are related to mutations of TFs [52]. In this study, we have identified 92 TFs as the regulator of the RPL30. Rahman MH et al. have identified a number of T2D-associated TFs, including GATA2, E2F1, FOXC1, FOXL1, YY1, NFIC, NFYA, USF2, TP53, POU2F2, CREB1 [53], consistent with our study. MiRNAs can regulate gene expression at the post-transcriptional level by degrading or inhibiting the translation of target genes [52]. Besides, E2F1, TP53, YBX1, EGR1, CTCF, AR and CREB1 was identified to play pathogenic roles in the occurrence and progression of IPF [54]. In addition, we have obtained 54 miRNAs of RPL30. Among these identified miRNAs, miR-3680-3p was reported may play important regulatory roles in disease mechanisms of T2DM with comorbid Tuberculosis [55]. Furthermore, the plasma expression signature of miRNA-126 and miRNA-320 are down-regulated in T2D, which was able to predict the further development of the disease in 70% cases [56]. Another study indicated that miRNA-557 and miRNA-17-3p were the potential therapeutic targets in IPF [54].

As far as we know, this is the first study to detect the role of shared genes in T2D and IPF through bioinformatics analysis. These results provide useful information that enhances our understanding of the pathophysiological process of T2D and IPF, and provide potential diagnostic and therapeutic targets for T2D and IPF. Nevertheless, this research study also has limitations. First, RPL30 has not been validated in experiments. Second, RPL30 was validated in patients with T2D or IPF, not with both diseases. The lack of a dataset including patients with both T2D and IPF made the validation unattainable at the present time. Third, the TFs-gene-miRNA interaction networks were only based on predictions from public databases but not further validation.

## Conclusions

This bioinformatic study elucidates RPL30 as common hub gene in T2D and IPF. Importantly, RPL30 may have value as biomarkers for use in diagnosing and monitoring T2D and IPF. Meanwhile, 92 TFs and 54 miRNAs were identified here that may be involved in regulation of RPL30. These results provide a new experimental foundation on which to further explore pathogenic mechanisms associated with T2D and IPF. Nevertheless, the specific mechanisms of RPL30 in T2D and IPF require further investigation.

## Data availability statement

The original contributions presented in the study are included in the article/Supplementary Material. Further inquiries can be directed to the corresponding authors.

## Supplemental Material

**Supplementary Figure S1.**
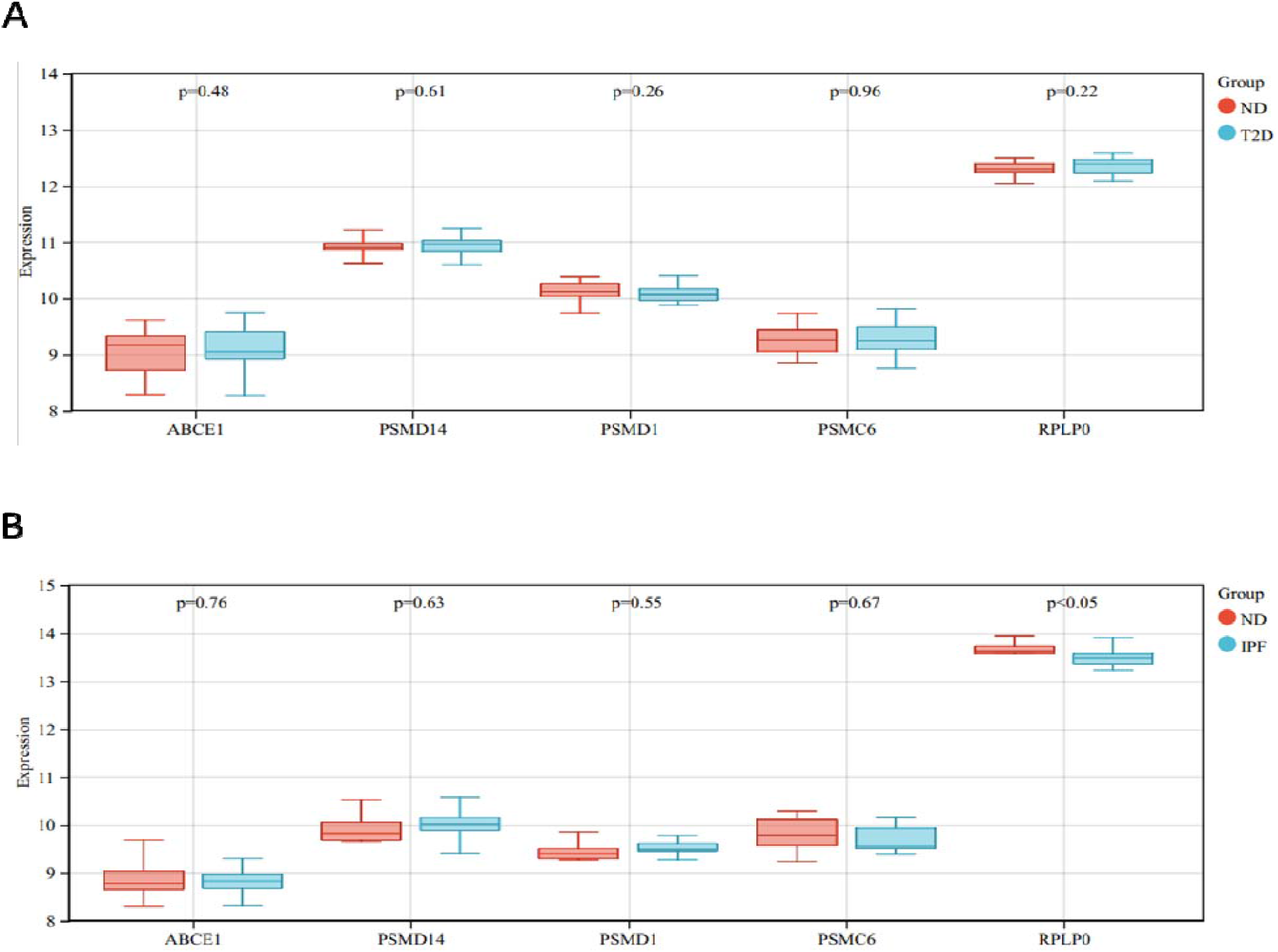
Validation of hub genes. (A) Hug genes were validated in GSE166467. (B) Hug genes were validated in GSE24206.

